# Numerical simulation of the viral entry into a cell driven by the receptor diffusion

**DOI:** 10.1101/822015

**Authors:** T. Wiegold, S. Klinge, R. P. Gilbert, G. A. Holzapfel

## Abstract

This study focuses on the receptor driven endocytosis typical of viral entry into a cell. A locally increased density of receptors at the time of contact between the cell and the virus is necessary in this case. The virus is considered as a substrate with fixed receptors on its surface, while the receptors of the host cell are free to move over its membrane, allowing a local change in their concentration. In the contact zone the membrane inflects and forms an envelope around the virus. The created vesicle imports its cargo into the cell. The described process is simulated by the diffusion equation accompanied by two boundary conditions. The first boundary condition states that the conservation of binders expressed as the local rate of change of density has to be equal to the negative of the local flux divergence. The second boundary condition represents the energy balance condition with contributions due to the binding of receptors, the free energy of the membrane, its curvature and the kinetic energy due to the motion of the front. The described moving boundary problem in terms of the binder density and the velocity of the adhesion front is well posed and relies on biomechanically motivated assumptions. The problem is numerically solved by using the finite difference method, and the illustrative examples have been chosen to show the influence of the mobility of the receptors and of their initial densities on the velocity of the process.

**SIGNIFICANCE:** The receptor driven endocytosis represents one of the most important mechanisms for the viral entry into a cell. However, the high velocities and small characteristic length scale cause many difficulties during the experimental investigation of such a process. This calls upon the application of virtual computer simulations investigating the process parameters and identifying factors inhibiting or completely ceasing the viral entry into cells. The development of methods for the optimization of the cell immunity system is aimed to as the final goal.

## INTRODUCTION

The intense study of cell mechanisms has provided an important insight into the uptake of various substances into a cell including viruses. Viruses cover a broad variety of shapes and sizes. The most common are sizes that range between tens to hundreds of nanometers (1, 2). Many investigations of cellular processes are limited due to the nanoscale. However, the progress in technological fields such as the electron microscopy has recently led to new discoveries and a deeper understanding on cell activities. In addition, numerical methods in biomechanics and biomathematics are now of more importance. With increasing hardware capabilities more complex and computationally expensive models are possible. The broad spectrum of numerical methods includes various applications in biomechanics. Some examples are the multiscale modeling of materials such as cancellous bone (3, 4), the modeling of cell membranes with structural elements such as shells (5, 6), or the simulation of the uptake of specific substances through the cell membrane, the cytoplasm, into the cell nucleus (7, 8).

Medical investigation on the basis of nanotechnology dates back to 1965 describing examples of lipid vesicles (9), later known as liposomes. Nowadays, nanotechnology has become more and more present in drug delivery systems, covering polymeric micelles, quantum dots, liposomes and many more. These systems provide opportunities to target specific cells or the control of drug release rates (10) and enable multistage delivery systems in a time-controlled fashion (11). They find use in many applications such as gene delivery, tissue engineering and tumor destruction (12).

In nature, a biological cell is surrounded by a plasma membrane, which acts as the interface between the cell and its surrounding environment. However, the membrane is not absolutely impermeable and a transport of particles through the membrane is still possible. Among many different mechanisms, the most common process for this purpose is the so-called endocytosis (13). The main focus of the investigation of endocytosis has been on clathrin-mediated endocytosis (CME). A greater understanding of the process has become increasingly present. During the CME, proteins create clathrin-coated pits which eventually build whole vesicles (14, 15). Similar processes to endocytosis are phagocytosis, responsible for the uptake of larger particles and macropinocytosis, responsible for the uptake of fluids. Both processes are caused by actin-dependent mechanisms (16, 17).

This work deals with the modeling of the endocytosis process in order to simulate the viral entry driven by receptor diffusion. In different chemical and biochemical contexts, the receptor binding has been extensively studied experimentally (18, 19) and theoretically (20–23). As described in (24) and (25), the binding of viral ligands is the main mechanism which leads to entry by fusion.

The most recent approaches are kinetic models for ligand-receptor binding. A mathematical framework to model viral entry in general is provided in (26). Therein, two alternative mechanisms driving the receptor binding process, a random and a sequential one, are introduced. The presented framework shows that the viral entry process is independent from the virus except for its receptor density.

The qualitative features of the viral entry process are dictated by the cell properties, particularly by the mobility of all receptors. The influence of this factor is discussed in (27). Based on the idea that the adhesion bond evolves along an admissible path, the mobility controls the adhesion bond rate. Several analytical models for this problem have already been established, see, for example, (27). However, these models still have some drawbacks, such as their dependence on several assumptions. These assumptions are motivated more mathematically than biomechanically. In the following, an approach is chosen, which focuses on the energetic aspects during the endocytosis process in order to provide a well-posed fully biomechanically motivated description.

This paper is structured as follows: The section ‘Methods’ gives a general overview of the uptake process followed by a recapitulation of the main aspects of the free energy characteristic for the endocytosis process. This section also focuses on the definition of boundary conditions accompanying the driving diffusion differential equation. The boundary conditions deal with the flux balance and the energy balance at the adhesion front. The numerical implementation is discussed in the last part of the section ‘Methods’. To this end the finite difference method is applied. The section ‘Results and Discussion’ deals with the representative numerical examples simulating the viral entry into a cell. Particular attention is paid to the influence of the mobility of the receptors as well as to the influence of different initial density configurations on the velocity of the process. The effect of the receptor cooperativity is also studied. The paper finishes with a conclusion and an outlook.

## METHODS

### Description of the uptake process

In a basic view of mechanical adhesive contact between elastic surfaces, two phenomena having a considerable influence on the underlying process counteract each other. A reduction in the free energy when surfaces with bonding potential come into contact benefits the process, whereas an increase in free energy due to elastic deformation required to fit their shapes aggravates the process. In the classic Hertzian theory of elastic deformation (28), two bodies coming into contact deform in the contact area in such a way that they perfectly fit. According to this approach any surface interactions such as Van der Waals forces, which are induced by charge polarization in electrically neutral molecules in close proximity, are excluded. However, these ‘non-material’ effects have a significant influence on the direct contact interaction. This is illustrated by considering small elastic objects consisting of crystalline materials processed in a controlled environment. Such crystals show the appearance of unfulfilled or dangling chemical bonds distributed over a free surface. Bringing such objects into contact reduces the free energy of the system by forming bonds between the two surfaces. The objects joined in this way will not separate without additional work. Hence, not only compressive traction due to bulk elasticity, but also an adhesive or tensile traction contributes to the contact. The same effects, both attractive and resisting interactions, appear in adhesive contact of biological cells. However, due to their characteristic properties compared to engineering materials, significant differences occur in this case. Having a remarkably lower elastic modulus than engineering materials weakens the influence of the effect of elastic energy variations during contact. Furthermore, cells are characterized by a fluid-like in-plane behavior. This enables the receptors of the cell to move within its membrane, enabling new methods of incorporating free energy variations in the modeling of adhesive contact.

In order to depict the process of viral entry into a cell, the situation presented in Fig. 1 is considered. This simplified case assumes rotational symmetry which corresponds to a spherical virus and homogeneous distribution of receptors on both surfaces (Fig. 1(a)).

**Figure 1:**
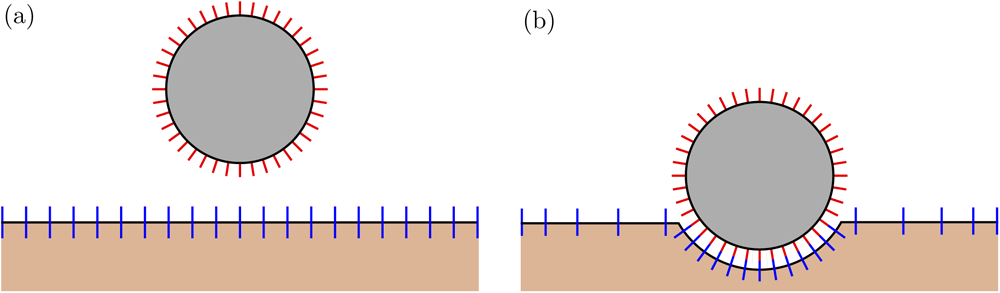
(a) Initial configuration of the cell surface and the virus in a 2D setup; (b) state during the uptake where the virus is partially connected to the cell.

Of course, in order to consider a more general case an extension is straightforward. In the initial state, the virus has not yet reached the cell surface (Fig. 1(a)). Upon first contact, the virus gradually connects to the cell (Fig. 1(b)). In order to establish a connection between the virus and the cell, a generic repulsion between their surfaces needs to be overcome. The connection by binding receptors of the virus to receptors of the cell reduces the internal energy of the system. Upon completing a single receptor-ligand bond the internal energy is reduced by *k T C*_b_, where *k* is the Boltzmann constant, *T* the absolute temperature and *C*_b_ a constant factor.

The quantity driving the uptake process is the receptor density ξ. Initially, the density of receptors on the cell surface amount to ξ_0_ and the corresponding counterpart, the receptor density on the virus surface, amounts to ξ_eq_. In general, it holds that the density of receptors on the virus is larger than the one on the cell surface and that the virus receptors are fixed, whereas the receptors of the cell are free to move across the membrane. Upon contact, receptors of the cell diffuse over the surface, connect to the receptors of the virus and build an envelope around the virus. At the end of the process the envelope is closed over the virus which has fully entered the cell.

As opposed to metallic or covalent bonds, the bonds created during biological adhesion are relatively weak. Since the cell receptor density is the lower one, in general, it dictates the amount of reduction in the internal energy. Typically, the resisting potential due to generic repulsion exceeds the reduction in internal energy of the initial configuration of the system for a unit area of the membrane at ξ_0_. Therefore, additional influences facilitate the creation or dissolving of chemical bonds. Possible influences are catalytic agents, small temperature changes and small mechanical forces. It appears, that a local change in receptor density is necessary in order to create an adhesion zone between the virus and the cell. An increasing local receptor density results in a greater reduction in the free energy by completion of each additional bond. When the cell receptors and the virus receptors are close to each other, a permanent interaction due to thermal stimulation is present.

In the framework of chemical rate theory two distinct cases are differentiated (29). The two cases are dictated by the relation between the virus receptor density and the cell receptor density. In the area where ξ < ξ_eq_ holds, the rate of bond breaking exceeds the rate of bond forming, so that no adhesive contact is established. In the area where ξ > ξ_eq_ holds, the rate of bond forming exceeds the rate of bond breaking, and adhesive contact is established. For the limit case ξ = ξ_eq_ a state of chemical equilibrium of the bonding reaction is achieved. Adhesion is assumed to occur locally in the region where the receptor density matches the equilibrium state and stays on this value throughout the process.

As an illustration, a schematic distribution of receptors over the cell and the virus is depicted in Fig. 2(a), whereas the corresponding density profile is shown in Fig. 2(b). In the adhesive zone, the receptor density is constant and amounts to ξ_eq_. Outside the adhesive zone the density grows and tends towards the initial receptor density of the cell ξ_0_ far away from the contact area. Since the size of the cell is magnitudes larger than the virus, we assume the receptor density far away from the adhesion front to stay constant 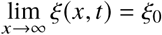. Consequently the flux *j* of receptors vanishes 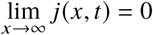.

**Figure 2:**
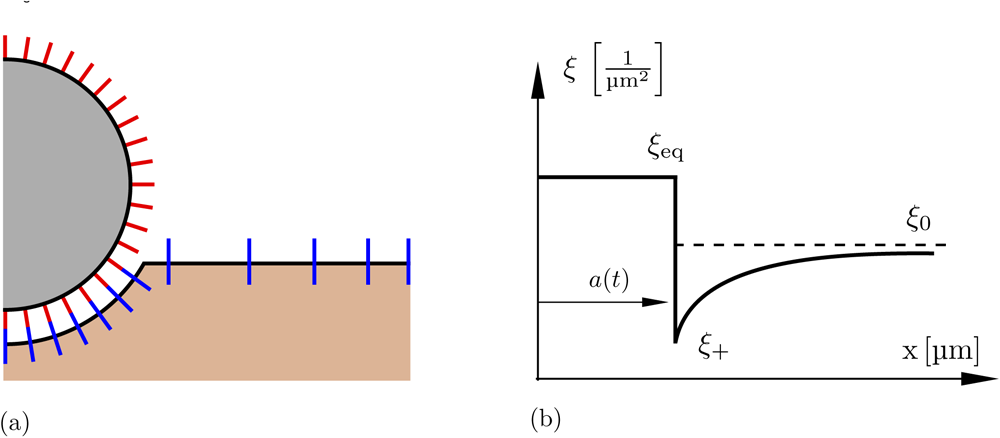
(a) Schematic diagram for the receptor distribution over the cell and virus surface; (b) typical density profile.

One more peculiarity of the density profile is the front of the adhesion zone, where a jump of receptor density occurs. The position of the front is depicted by a function of time *a*(*t*). The values typical for the adhesion front play an essential role in the model to be described, and are denoted by the subscript + in the subsequent text. For example, ξ_+_ denotes the receptor density at the front. No generation or destruction of receptors occurs during the process, such that their movement is only responsible for the change in density.

The previous explanation shows that the whole process is regulated by the diffusion of receptors over the cell surface and their gathering in the adhesion zone. Accordingly, the motion of the receptors will be described by the diffusion differential equation, i.e.

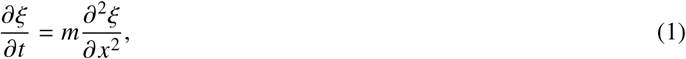

which defines the relation between the temporal and the spatial changes of the receptor density weighted by the mobility parameter *m*. Its evaluation gives insight into the evolution of receptor density for every point in front of the adhesion front *a*(*t*) < *x* < ∞. Following Fick’s first law, the receptor flux *j* is proportional to the gradient of density, i.e.

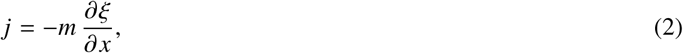

which leads to the alternative expression of the diffusion equation in terms of the receptor flux according to

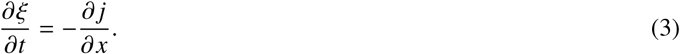

This relationship states that the change of receptor density in time has to be equal to the spatial change in the flux. Equation (1) is a partial differential equation of second order and requires two additional boundary conditions in order to determine the complete particular solution. These two conditions will be defined in the upcoming sections.

### Process characterization

In order to define the free energy characteristic of the simulated process, the system including a large number of receptors is treated analogously to the case of an ideal gas with a large number of non interacting particles *N*. In such a case, the entropy of a single particle, belonging to a system in equilibrium, is expressed by *k* ln[(*A*/Λ^2^)/(*c/N*))]. Here, *A* is the considered surface, *N/A* is the areal density ξ, *c* is a numerical factor and Λ a molecule length scale. However, later two quantities (*c* and Λ) do not play any role for the description of our process since it does not depend on the absolute entropy but on its change. Assuming the initial state of the cell surface as the reference state, the relative entropy at density ξ is calculated as follows (27)

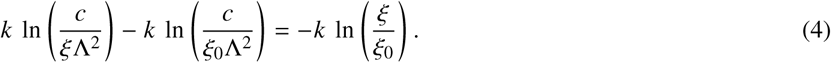

With this expression at hand, the free energy *E*_e_ per unit area of membrane surface associated to the receptor distribution at absolute temperature *T* turns into

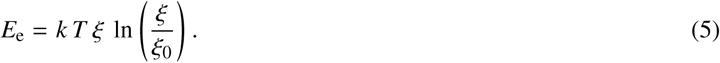

Moreover, the chemical potential χ is defined as the local change in the free energy per receptor,

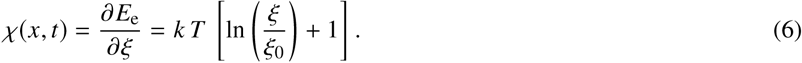

Finally, the mean receptor speed is assumed to be proportional to the spatial gradient of the chemical potential, i.e.

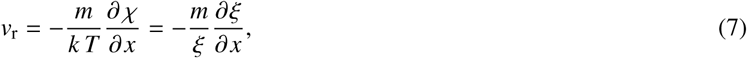

where the movement of the receptors is controlled by the mobility parameter *m*.

### Boundary conditions

The full description of the adhesion front motion relies on the differential equation (1) along with two boundary conditions: flux and energy balance.

### Flux balance

The first boundary condition is concerned with the quantitative description of the flux of receptors through the adhesive front. Following the Leibniz integration rule of the global form, this condition is derived from (3) as

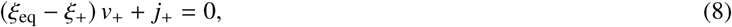

or by using Fick’s first law as

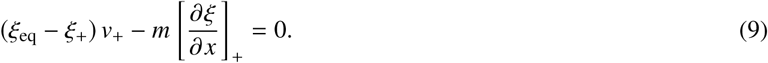

Here, the first term denotes the amount of receptors required for the forward movement of the front, and the second term denotes the amount of receptors provided by the flux. Equation (9) is consistent with the assumption (7) which can easily be shown as follows. First, the flux is assumed to be proportional to the receptor distribution ξ and the mean receptor velocity v_r_. Thus,

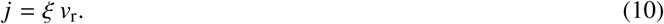

By incorporating Eq. (7) into Eq. (10), the flux turns into

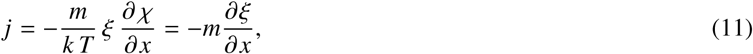

as predicted by Fick’s first law.

### Energy balance

The second boundary condition for the present moving boundary problem is provided by considering the energetic aspects of the front motion. The change of the receptor distribution as well as of the membrane shape lead to several contributions to the free energy of the system. However, the crucial observation is that the difference in the energy afore and behind the front results in the front movement, which is expressed as follows,

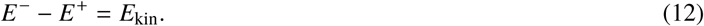

Here, *E*^−^ denotes the energy behind the adhesion front, *E*^+^ is the energy afore the front and *E*_kin_ is the kinetic energy of the front itself.

The term related to the energy behind the front is built of three contributions, all denoted by superscript −,

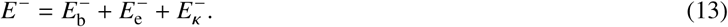

These terms have the following physical meaning: 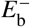 is the energy related to the binding of receptors, 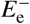 the energy related to the entropy and 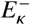 the energy related to the bending of the membrane. The reduction in the free energy due to the binding of receptors of the cell to receptors of the virus is defined as follows

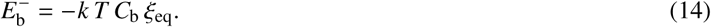

This term is proportional to the reduction of energy caused by a single bond between two receptors −*k T C*_b_, and to the total amount of created bonds ξ_eq_ dictated by the virus. The constant factor *C*_b_ typically takes values in the range 5 < *C*_b_ < 35. The second term describes the energy associated with the entropy of receptors

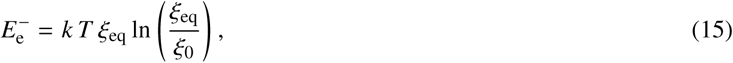

which is required to bring the density from its reference value ξ_0_ to the density of the virus ξ_eq_. This term will result in an increase to the free energy since it holds ξ_0_ < ξ_eq_. The third term of (13) is concerned with the bending of the membrane caused by the geometry of the virus

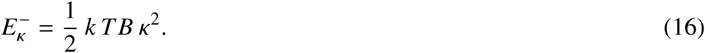

Here, a simplified case is considered corresponding to the theory of the bending of a plate. Factor *B* represents the bending stiffness, which is in the range of 10 to 30 and κ = 1/*R*_v_ represents the curvature, which is for a spherical virus constant and depends on the radius of the virus *R*_v_. Thus the whole energy behind the front is then defined by the expression

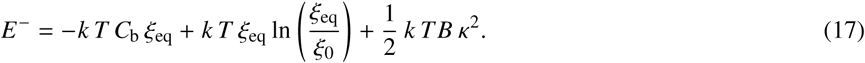

In the second step we consider the energy afore the front, denoted by superscript +. Binding between the cell and the virus exclusively takes place in the area behind the front and thus does not have any influence on the energy afore the front. However, corresponding parts 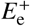, the energy related to the entropy and 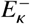, the energy related to the curvature of the membrane remain available. Moreover, a term 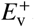, the energy related to the movement of receptors also has to be taken into consideration. In summary, the following terms can be counted afore the front:

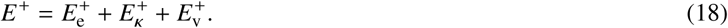

In the present contribution, we assume that the curvature behind the front is much smaller than the one caused by the contact with the virus. This justifies the assumption of a vanishing influence to the energy associated to the bending of the membrane

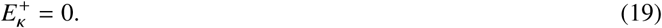

The energy afore the front related to the entropy is expressed in the same way as the energy behind the front as

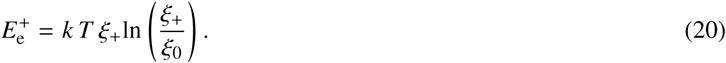

It describes the energy needed in order to bring the initial receptor density ξ_0_ to the value ξ_+_. Contrary to the contribution behind the front, this term results in a reduction of the free energy since ξ_0_ > ξ_+_. The contribution due to the movement of the receptors reads

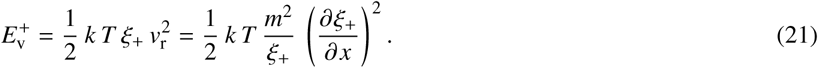

It represents the kinetic energy of all receptors afore the front moving towards the front with their corresponding velocity v_r_. With Eqs. (19) - (21), the total energy afore the front is defined by

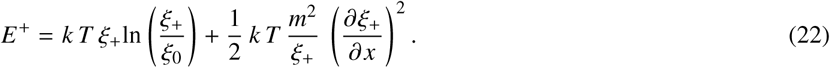

Finally, the difference between the energies of the two sides of the front acts as driving force for the front movement. The kinetic energy of the front is then characterized by the ‘mass’ of the front *k T* ξ_eq_ and the front velocity v_+_

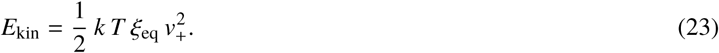

Combining Eq. (17), (22) and (23) leads to the expression

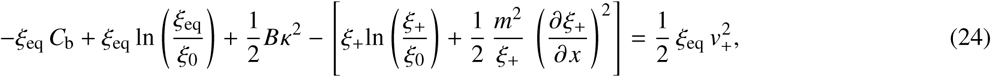

which is the final form of the second boundary condition and closes the formulation of the moving boundary problem.

### Implementation

In summary, the change of the receptor distribution is described by a system of differential equations consisting of (1), (9) and (24). The finite difference method has been chosen for the solution of the underlying system of differential equations. According to this approach, all derivatives are replaced by expressions dependent on discrete values of the function for the nodes of a chosen lattice. Thus the differential equations are transformed into a system of algebraic equations. An implicit scheme is used, with the following approximations for the derivatives

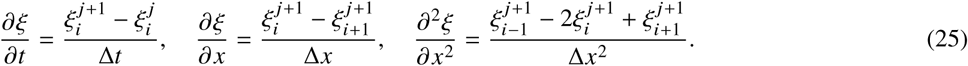

Here, subscript *i* denotes the spatial position and superscript *j* the time. The implementation of relationships (25) into the system (1), (9) and (24) leads to the following discretized formulation of the problem:

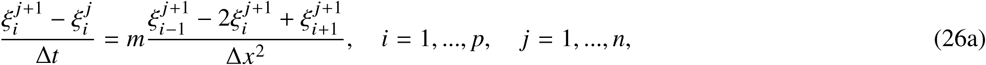

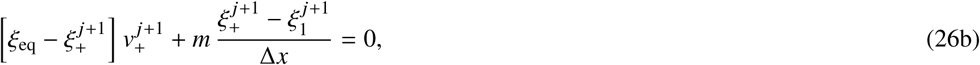

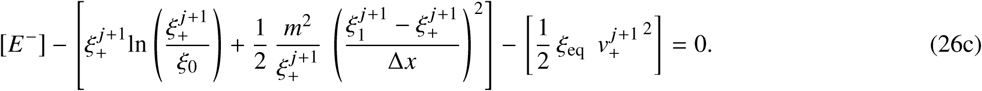

In Eq. (26a), the variable *p* refers to the total number of points afore the front, except the last point, where the influence of the flux vanishes, and where the receptor density is kept at the initial value ξ_0_. The variable *n* refers to the total number of time steps. Furthermore, Eqs. (26a) - (26c) are boundary conditions valid at the front. In Eq. (26c), *E*^−^ is an abbreviation for the part provided in (17). This term does not depend on the density ξ_*i*_ and quantities at the front ξ_+_ and v_+_, and thus represents a constant during the process.

## RESULTS AND DISCUSSION

The numerical examples chosen simulate the process of virus uptake into the cell. In the simulations, it is assumed that a spherical virus of size *D* = 0.05 µm comes into contact with a much larger cell such that the cell curvature is neglectable (Fig. 1). The initial density of cell receptors is ξ_0_ = 1000 µm^−2^, whereas the initial density of virus receptors is ξ_eq_ = 4800 µm^−2^. Time increment ∆*t* = 1*e*^−4^ s and space increment ∆*x* = 1*e*^−3^ µm are chosen for the numerical simulations. An overview of the process parameters is given in Table 1. The values belonging to the corresponding admissible ranges have been chosen.

**Table 1:** Process parameters used in the simulations.

| **Material parameters** |  |  |  |
| --- | --- | --- | --- |
| Receptor density on cell surface | $\xi_0$ | 1000 | $\mu\text{m}^{-2}$ |
| Receptor density on virus | $\xi_{\text{eq}}$ | 4800 | $\mu\text{m}^{-2}$ |
| $C_b$-Parameter | $C_b$ | 5 | — |
| Bending stiffness coefficient | $B$ | 30 | — |
| Curvature of the virus | $\kappa$ | 40 | $\mu\text{m}^{-1}$ |
| Mobility parameter | $m$ | 0.5-1 | $\mu\text{m}^2/s$ |

The first group of simulations studies the change of the cell receptor density during the process and the front motion. The density profiles for different time steps during the simulation are presented in Fig. 3. The diagrams show a fast decrease in receptor density, and at the front it decreases fast at the beginning of the process. After 300 time steps, the density at the front only amounts to ≈ 50% of its initial value. This rapid decline of the density at the front slows down in the course of the further process.

**Figure 3:**
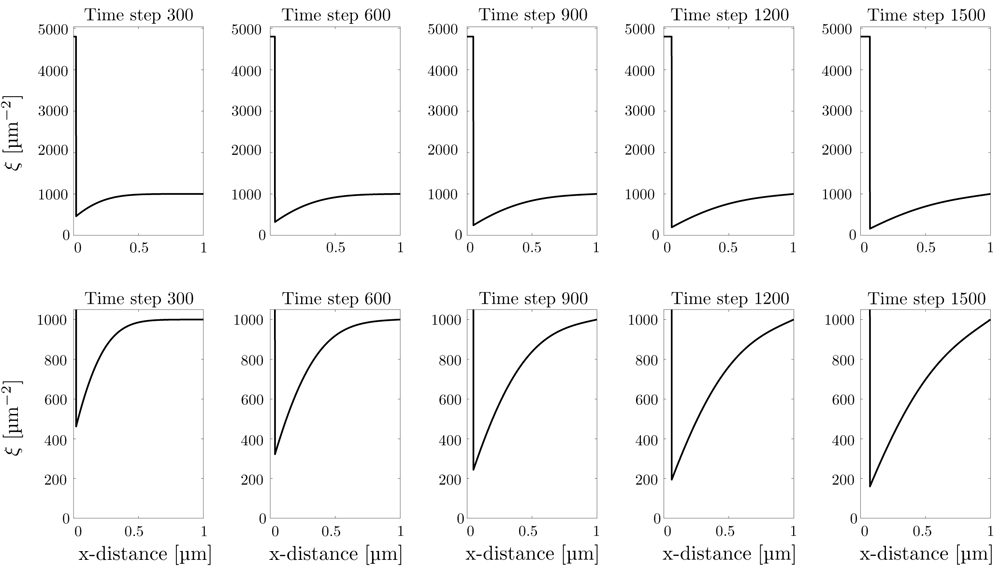
Receptor density ξ over the cell surface *x* for the first 0.15 s. The upper row shows the complete density profile whereas the lower row shows a zoomed in state, allowing a more precise visualization.

Figure 4 visualizes the advancement of the front and the position of the virus during its entry into the cell. The position of the virus is related to the position of the front through the length *a*, determining the size of contact area. Due to the symmetry of the chosen example, a direct visualization of the vesicle in 3D is possible (Fig. 5). The increasing number of time steps between four states indicates the gradual decrease and final stagnation of the velocity of the process, an issue also studied in the following example.

**Figure 4:**
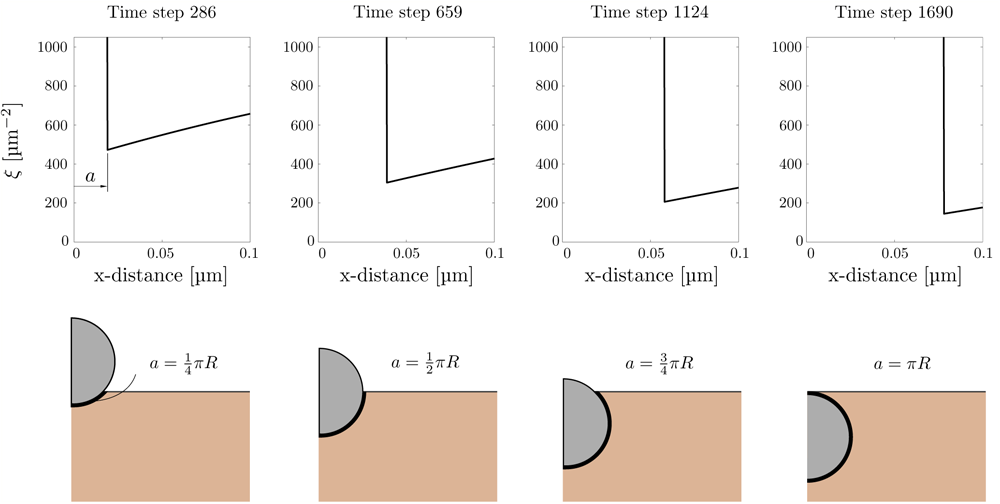
Visualization of the front motion and of the formation of the envelope around the virus with the diameter *D* = 0.05 µm.

**Figure 5:**
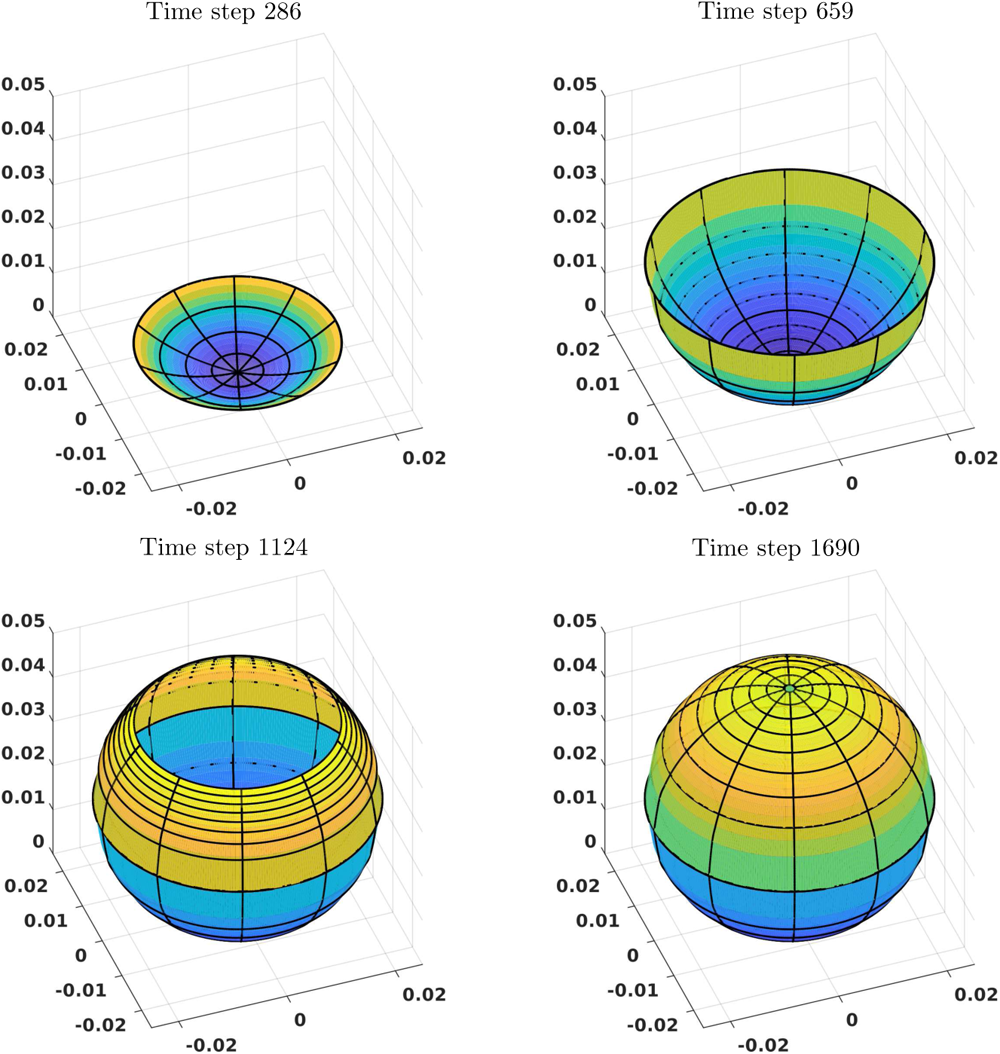
3D-Visualization: the formation of the envelope over the virus with the diameter *D* = 0.05 µm. Snapshots for the time steps 286, 659, 1124, 1690.

The governing equation (1) of the process depends on a single process parameter, namely on mobility *m*. The parameter represents a measure for the capability of receptors to move over the cell surface, and thus is in a direct correlation with the amount of receptors provided for the adhesion with the virus. The influence of the mobility on the velocity of the front and on the receptor density has been studied on the basis of a set of simulations, as shown in Fig. 6. Here, the mobility parameter *m* has been varied in the range 0.5 µm^2^/s-1 µm^2^/s. Figure 6(a) shows the dependence of the velocity v_+_ on the mobility and clearly approves the fast decrease of the velocity at the beginning followed by a stagnation, already observed in the previous test (Fig. 4). The value of the mobility does not affect the form of the velocity diagrams. However, as expected, a higher velocity corresponds to a higher mobility. This observation is in agreement with the physical character of the mobility describing the ability of receptors to move towards the adhesion zone. For lower values of *m*, fewer receptors are provided to connect the cell with the virus. Therefore, the evolution of the adhesion zone and the velocity of the front are slowed down. An analogous trend is observed for the dependency of the receptor density at the front on the mobility (Fig. 6(b)).

**Figure 6:**
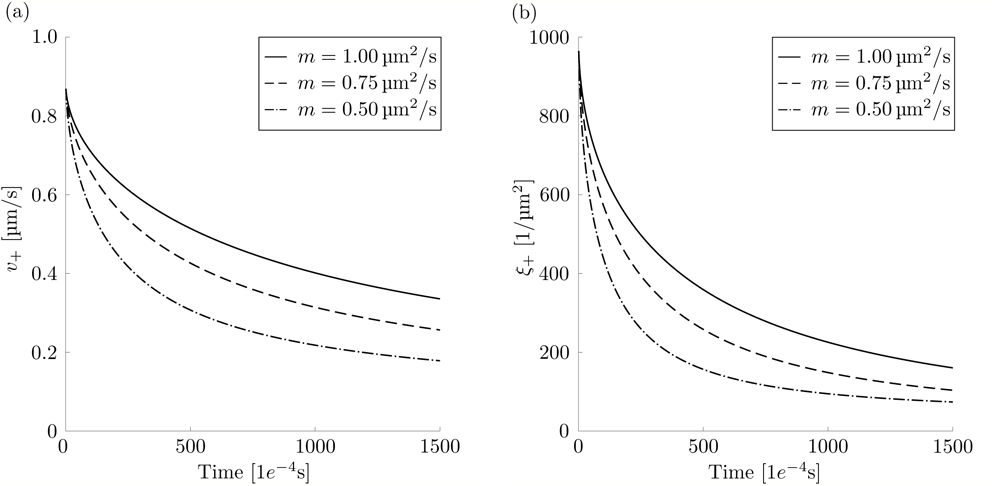
(a) Velocity of the adhesion front vs. time. (b) Evolution of the receptor density at the adhesion front. Mobility is varied in the range 0.5 µm^2^/s − 1 µm^2^/s.

An important influence on the process is also imposed by the fixed receptor density ξ_eq_ of the virus, which dictates the amount of receptors required for the virus-cell connection. The velocity of the adhesion front v_+_ for different values of ξ_eq_ is shown in Fig. 7(a). Here, the receptor density of the cell is set to ξ_0_ = 1000 µm^−2^. The form of the velocity diagrams does not change, although the different constellations are taken into consideration. The velocity of the adhesion front v_+_ decreases with increasing density ξ_eq_, which is to be expected since a larger number of receptors is necessary in order to achieve a front advancement. Similar simulations are conducted for different values of the initial receptor density ξ_0_, while the receptor density of the virus is set to ξ_eq_ = 4800 µm^−2^ (Fig. 7(b)). Again, the initial configuration does not affect the form of the diagrams, while a larger density ξ_0_ corresponds to higher velocities. The required amount of connected receptors has been fixed at a constant value in all the simulations performed. However, only the initiation of the process requires a higher amount of bonds. Once contact between the cell and the virus is established, the number of necessary receptors decreases. The amount of bonds required for the contact between the cell and the virus cannot fall below a minimum value. The evolution of the required cell receptor density can be easily implemented in the developed code by assuming ξ_eq_ to be a function of time. The simulations in this case (results not shown here) indicate an accelerated viral entry into the cell as a consequence of the decrease of the required receptor density.

**Figure 7:**
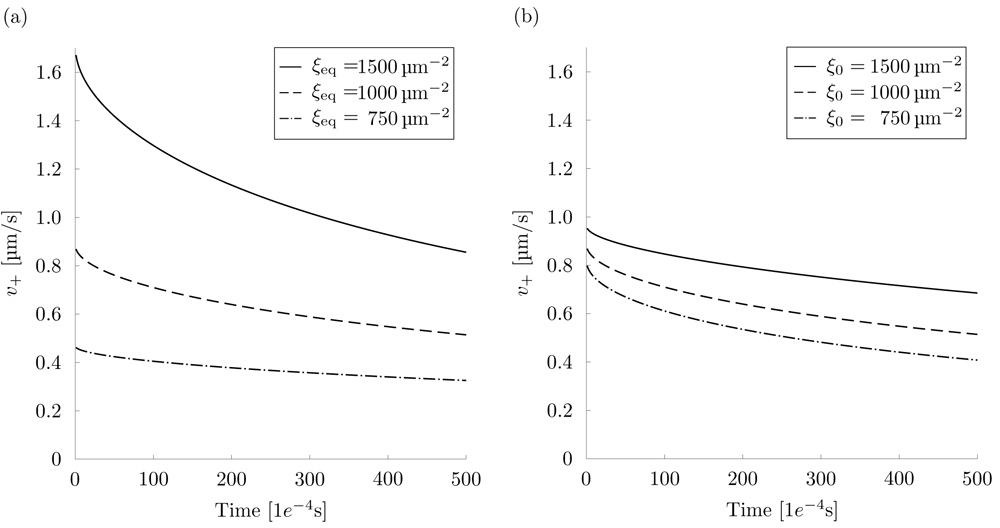
(a) Evolution of the velocity of the adhesion front for different densities ξ_eq_. Density ξ_0_ is set to 1000 µm^−2^. (b) Evolution of the velocity at the adhesion front front for different initial densities ξ_0_. Density ξ_eq_ is set to 4800 µm^−2^.

### Cooperativity

Among others, cell adhesion deals with cooperativity, an effect which is explained by considering a patch of unit length (Fig. 8). As soon as receptors create bonds they smooth out the surrounding membrane which makes it easier for additional receptors to create a bond and strengthens the adhesion between the virus and the cell membrane (30, 31). This effect is known as cooperativity. It has extensively been investigated experimentally and theoretically. Different experiments are performed depending on the state of the adhesion process. The fluorescence recovery experiments are performed in order to analyze the equilibrated contact zone during the process, whereas the micropipette experiments are performed in order to analyze the initial contact. Lipid vesicles with anchored receptor molecules are often used in order to resemble important aspects of cell adhesion. In order to study the binding cooperativity, two classes of numerical models are considered. The first class describes the membranes as continuous in space with continuous concentration profiles on the membrane (32). The second class describes the membranes discrete and the receptors as single molecules (33). Numerical solutions of the dynamic properties are studied by reaction-diffusion equations in the first class (34) of models and by Monte Carlo simulations in the second class (35). The information obtained in such a way is complementary to the model presented in this contribution.

**Figure 8:**
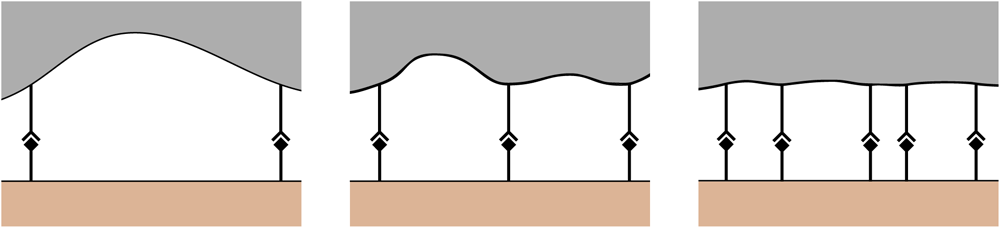
Schematic representation of cooperativity during endocytosis (the free receptors are not shown).

The cooperativity reduces the total amount of required receptor bonds in order to connect the virus with the cell according to

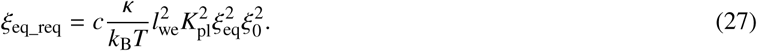

Here, ξ_eq_req_ is the required amount of receptors that need to bind in order to create adhesion between the virus and the cell. *c* is a dimensionless prefactor acquired from Monte Carlo simulations usually in the range 10 - 15. The quantity *l*_we_ is the binding range depending on the interaction range of the two binding sites, of the flexibility of their molecules and of the membrane anchoring. It describes the difference between the smallest and the largest local membrane separation at which the receptors can bind. Quantity *K*_pl_ is the two-dimensional equilibrium constant in the case of two opposing planar, supported membranes within binding separation of the receptor–ligand bonds.

Two examples are given in order to analyze the influence of cooperativity and different binding ranges. The first group of simulations considers a virus with the lower half covered by receptors with a smaller binding range and the upper half by receptors with a larger binding range. The lower half is characterized by a binding range of *l*_we_ = 1 nm resulting in the required receptor density ξ_eq_req_ = 2265 µm^−2^, while the upper half is characterized by a binding range of *l*_we_ = 1.2 nm resulting in the required receptor density ξ_eq_req_ = 3262 µm^−2^. An opposite situation is considered in the second group of simulations (Fig. 9). Numerical results for the described examples are shown in Fig. 10. The transition between the areas with different receptor types manifests itself by either a jump or a kink in the corresponding diagrams. The velocity is affected most by the change of the required density. In the area with a smaller binding range less receptors are required, which significantly increases the velocity of the process. The diagrams for the second setup show similar results to the first setup, however, the change from the lower to the upper half is significantly delayed.

**Figure 9:**
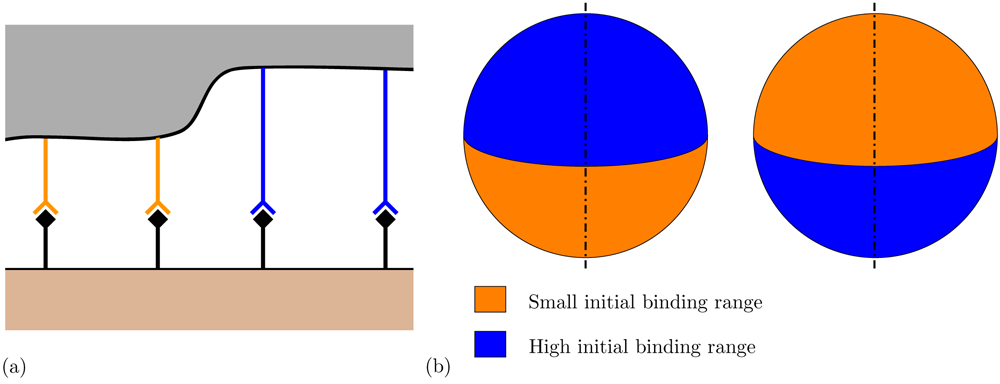
(a) Contact with the virus with two kinds of receptors; (b) spatial distribution of different types of receptors on the virus membrane for two chosen configurations.

**Figure 10:**
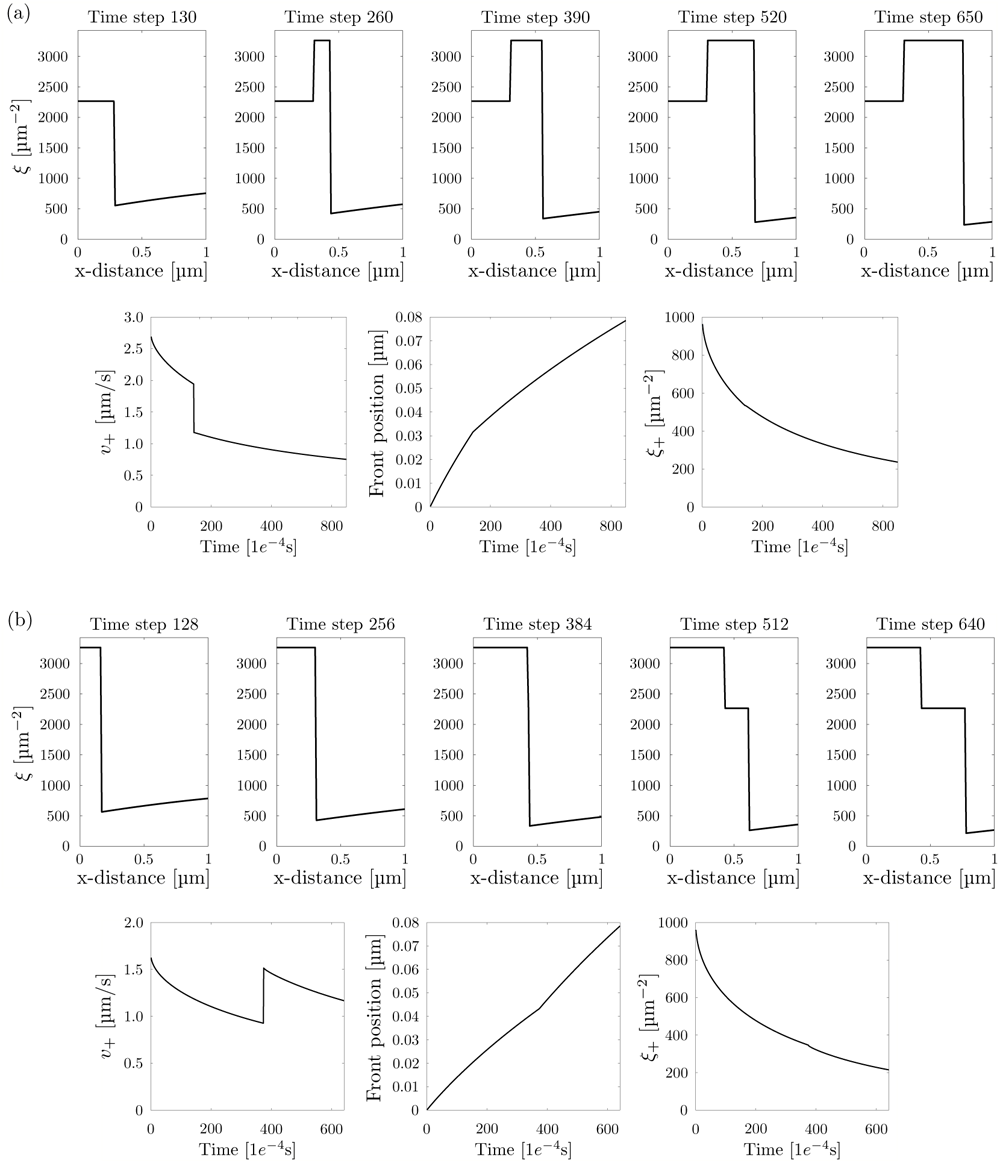
(a) Influence of the cooperativity for a smaller binding range in the lower half ξ_eq_req_ = 2265 µm^−2^ and a larger one in the upper half ξ_eq_req_ = 3262 µm^−2^; (b) Influence of the cooperativity for a larger binding range in the lower half ξ_eq_req_ = 3262 µm^−2^ and a smaller one in the upper half ξ_eq_req_ = 2265 µm^−2^ Top row: Receptor density over the cell surface for different time steps. Bottom row: Velocity of the front, position of the front and receptor density at the front over time. Chosen process parameters are *K*_pl_ = 0.55 *e*^−3^ and *c* = 13.

3D simulations of the viral entry for both chosen configurations are presented in Fig. 11. Here, the initial velocity is much higher in the first case such that the virus is almost enclosed at the time step 400. Contrary to this, the velocity at the end of the process is higher in the second case. Consequently, both viruses need approximately 600 time steps for their entry into the cell. Exact values are 611 and 622 time steps for the first and second example respectively. The values do not match exactly due to the different velocities at the beginning of the process and due to the transition between regions with different receptors.

**Figure 11:**
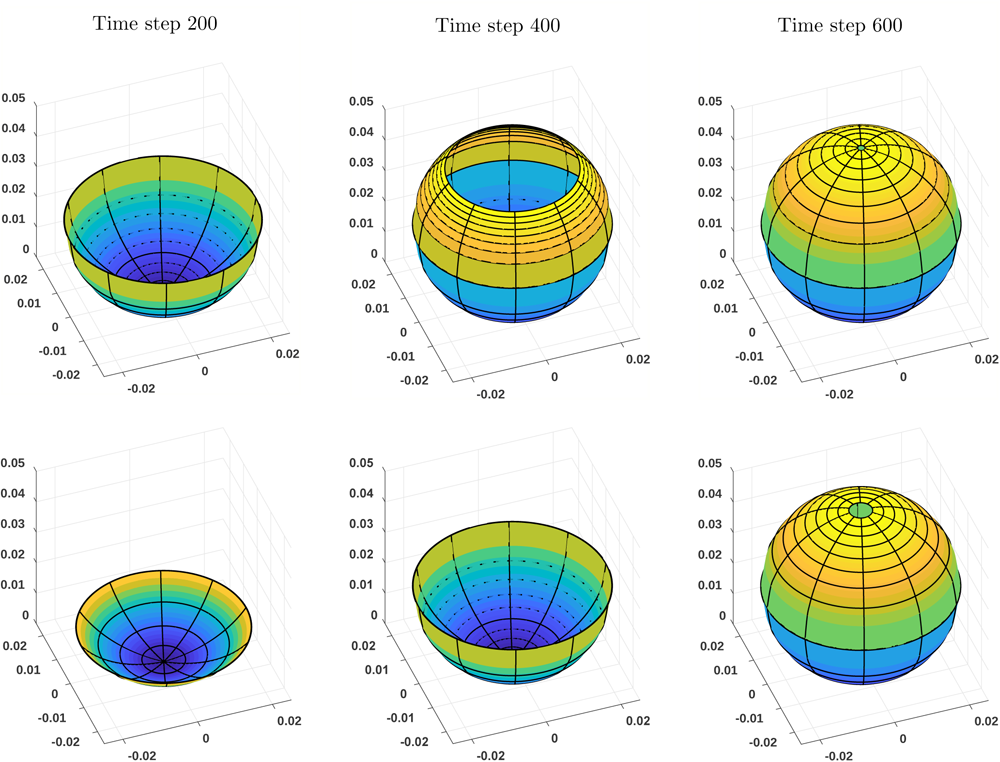
3D representation of the vesicle for two setups after 200, 400 and 600 time steps. Top row: Smaller required receptor density in the lower half (ξ_eq_req_ = 2265 µm^−2^), larger one in the upper half (ξ_eq_req_ = 3262 µm^−2^). Bottom row: Results for the setup with exchanged required receptor density. Process parameters are listed in the caption of Fig. 10.

## CONCLUSION

The present study focuses on the investigation of the viral entry driven by the receptor diffusion using the finite difference method as simulation technique. An approach based on the consideration of the energetic aspects yields a formulation avoiding ad-hock assumptions and simplifications. The movement of the receptors is described by a diffusion differential equation accompanied by two boundary conditions. The first boundary condition deals with the flux balance at the adhesion front, and the second boundary condition deals with the energy balance at the adhesion front.

The proposed model enables an efficient numerical simulation of the viral uptake into a cell. The obtained results show a fast change in the receptor density at the beginning of the process which then slows down as the time progresses. Characteristic parameters dictating the evolution of the receptor density are the mobility parameter and the initial density distribution of the virus and the cell. The study of their influence reveals a strong correlation between these parameters and the speed of the process. As expected, higher values for the mobility and lower ratios between the initial virus and the cell density result in a faster process procedure. Higher values for the receptor density of the virus result in a slow down of the process procedure. An important advantage of the model proposed is its compatibility with medical investigations, as shown in examples simulating the cooperativity. This effect describing a reduction in the required receptor density for bonding is related to the observations of fluorescence recovery and micropipette experiments.

The results presented in this work are concerned with a spherical virus penetrating a flat cell surface, which enables the taking of advantage of the symmetry and perform simulations in a two dimensional setup. However, an extension to a three dimensional setup has to be taken into account in order to analyze the receptor distribution for a non-spherical virus or a non-homogeneous receptor density of the cell. Furthermore, additional contributions, for example, caused by bending of the cell afore the front, can be considered in the energetic approach of the second boundary condition. Moreover, alternative expressions for bending lipid bilayers can be introduced in order to carry out even more realistic simulations.

## AUTHOR CONTRIBUTIONS

T.W., S.K., R.P.G. and G.A.H. equally participated in the development and verification of the model. T.W. carried out numerical simulations.

## ACKNOWLEDGMENTS

We gratefully acknowledge the financial support of the German Research Foundation (DFG), research grant No. KL 2678/7-1, and the Austrian Science Fund (FWF), research grant No. I 3431-N32. We also thank Dr. Matias Hernandez from the Max Planck Institute of Molecular Physiology at Dortmund, Germany for valuable discussions.

